# Chemical diversity in some biofouling organisms from the western coastal waters of Sri Lanka

**DOI:** 10.1101/2022.10.21.513247

**Authors:** R L Weerasinghe, R R M K P Ranatunga, S D M Chinthaka

## Abstract

Sri Lanka occupies a strategic position in the Indian Ocean, making the surrounding ocean one of the busiest in the region. The lack of fundamental studies has created a void regarding the physical and chemical behaviour of the fouling community. A few studies have been conducted to assess the subtidal biofouling communities and invasive threats in key ports and surrounding coastal waters. This study explores the chemical diversity and environmental resilience of nine marine macrofouling organisms through secondary metabolite-induced impacts on biofilm formation and volatile component analysis. The anti-settlement assay revealed that *Schizoporella errata, Botrylloides violaceus, Callyspongia diffusa*, and *Acanthella cavernosa* showed significant resistance against *Escherichia coli* settlement within the first 12 h (OD_600_ < 0.1). The identification of known compounds with a higher degree of antimicrobial activity, such as dodecanoic acid, methyl palmitate, β-caryophyllene and β-asarone, further supports the findings of anti-settlement activity of macrofouling organisms and likely plays a role in environmental resilience.

## 1. INTRODUCTION

Marine biofouling is a natural phenomenon that threatens an array of submerged surfaces in marine environments. It is the undesirable accumulation of marine organisms through the settlement of micro and macro-organisms on favourable surfaces, often resulting from the deposition of organic matter. Prolonged exposure to water initiates the formation of biofilms on susceptible surfaces via a continuous process of surface conditioning and recruitment in which the conditioned surface is colonised by microfoulers followed by soft and hard macrofouling, respectively (Bekiari et al., 2015; Ferreira-Vançato et al., 2020; Kim et al., 2021; Menchaca et al., 2014). The intensity and variety of the fouling community are governed by several factors such as substratum, locality, season, climate, immersion period, competition, and predation (Callow and Callow, 2002), ensuring the development of the macrofouling community as a highly dynamic and complex process.

The macrofouling community comprises algae, soft-fouling types such as sponges, tunicates, soft corals, and hard foulers such as barnacles, tubeworms, and bryozoans with adaptations to facilitate their settlement in a new environment (Callow and Callow, 2002). Larval stages and macroalgal spores possess remarkable abilities to discern suitable submerged substrates before committing themselves to life as sessile individuals. Most submerged surfaces are subjected to constant colonisation pressure since their initial contact with water (Richmond & Seed, 1991; Wahl, 1989). Over the last few decades, the expansion of the human population has posited an increase in the number and reach of artificial-submerged structures in the marine environment (Todd et al., 2019). The substantial increase in long-distance maritime activities and industrial water-use structures has also added to the attachment surfaces, creating more favourable settlement sites for the biofoulers in turbulent habitat settings (Ojala & Tenold, 2017; Wahl, 1989).

Changes in the hydrodynamics due to ship hull fouling have affected the economy negatively by increasing fuel consumption and biofouling management costs, in which the coating roughness and the additional weight of fouling contribute to the operational drag of vessels. The settlement of fouling organisms increases surface corrosion vulnerability, which inflates the maintenance cost excessively (Clarke Murray et al., 2013; Schultz, 2007; Schultz et al., 2010). In addition, biofouling is a known cause for introducing non-indigenous species (NIS) across borders, primarily by commercial shipping activities. Artificial static structures in ports often act as the first point of settlement for NIS, posing a biosecurity risk through the spread of invasive species (Drake & Lodge, 2007; Floerl et al., 2009; Hewitt et al., 2009; Sylvester et al., 2011). The negative impacts of biofouling can result in the rise of maintenance fees and incite production failures and safety concerns that necessitate the application of heightened management practices, which are unfavourable to the environment.

Sri Lanka is in a strategic position within the east-west international shipping route, surrounded by the Indian Ocean (Devendra & Muthucumarana, 2013), with constant exposure to biofouling pressure. The Colombo Port was ranked number 13th in the world in terms of facilitating mainline services according to the Drewry Port Connectivity Index (Ports & Terminals Insight, 2017). Thus, defining it as one of the busiest ports in the region, which requires constant monitoring to determine the presence of invasive species in its marine environs, a few studies have been conducted to assess the biofouling community in key ports and surrounding coastal waters of Sri Lanka through the deployment of artificial settlement structures by which access to subtidal fouling communities has been enabled (Jayasundara & Ranatunga, 2016; Marasinghe et al., 2016). Moreover, pilot projects such as Port Biological Baseline Surveys (PBBS) have provided a significant boost to biofouling research. However, the lack of fundamental studies has created a void regarding the physical and chemical behaviour of the fouling community, which hinders the practical application of biofouling control (Marasinghe et al., 2018; Marine Environment Protection Authority (MEPA); Ranatunga et al., 2015). One of the main objectives of this study was to understand the possible secondary metabolite-induced impacts on biofilm formation and development. Therefore, the experiment was designed to assess the chemical diversity and environmental resilience of macrofouling organisms with respect to their living habitats.

## 2. MATERIALS & METHODS

The study was carried out in Colombo Port and adjacent coastal waters to investigate the chemical potential of a selection of marine organisms. A pilot study was conducted to understand the community structure of fouling organisms in the port prior to selecting the sampling locations following Marasinghe et al. (2015) and Marasinghe (2019). The fouling community structure in the environment was observed before the selection of sampling sites for the study. Four locations with the port premises, Colombo International Container Terminal (CICT), Bandaranayake Quay (BQ), Passenger Jetty (PJ), and Unity Container Terminal (UCT), were selected for the study at the end (Figure 2.1).

**Figure 2.1.**
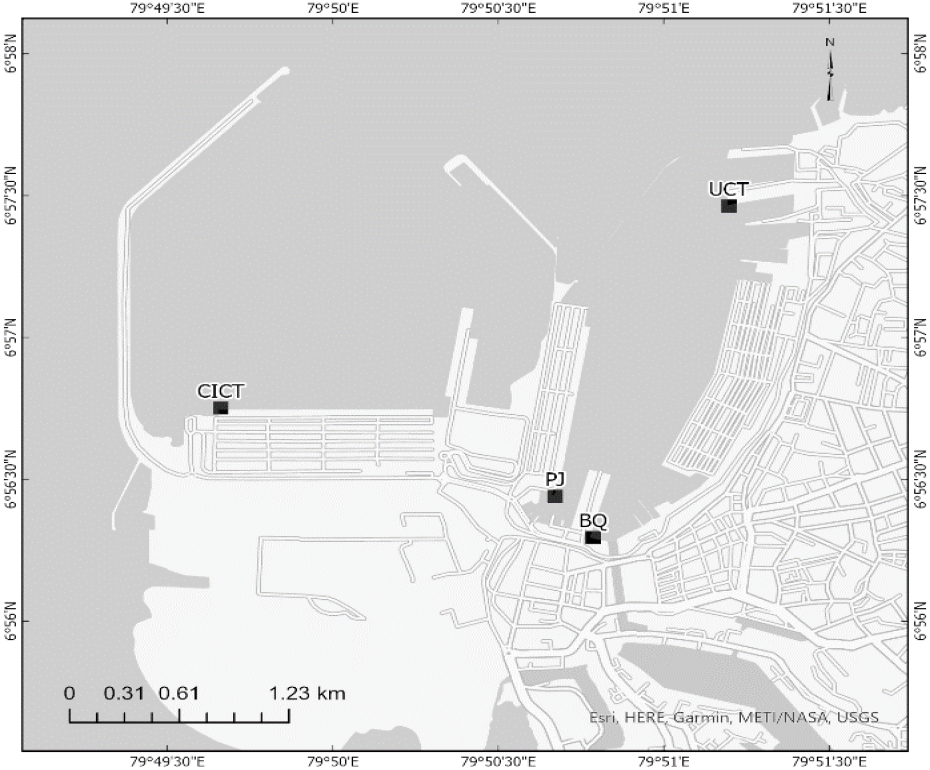
Sampling locations inside the Colombo Port: CICT, BQ, PJ, and UCT

In addition, two reference sampling locations, Kelani River Mouth (KLM) and Mount Lavinia Beach (ML), were selected outside of Colombo Port, and one reference location from the Vankalei reef off Mannar (M) of Sri Lanka (Figure 2.2 (a,b)).

**Figure 2.2:**
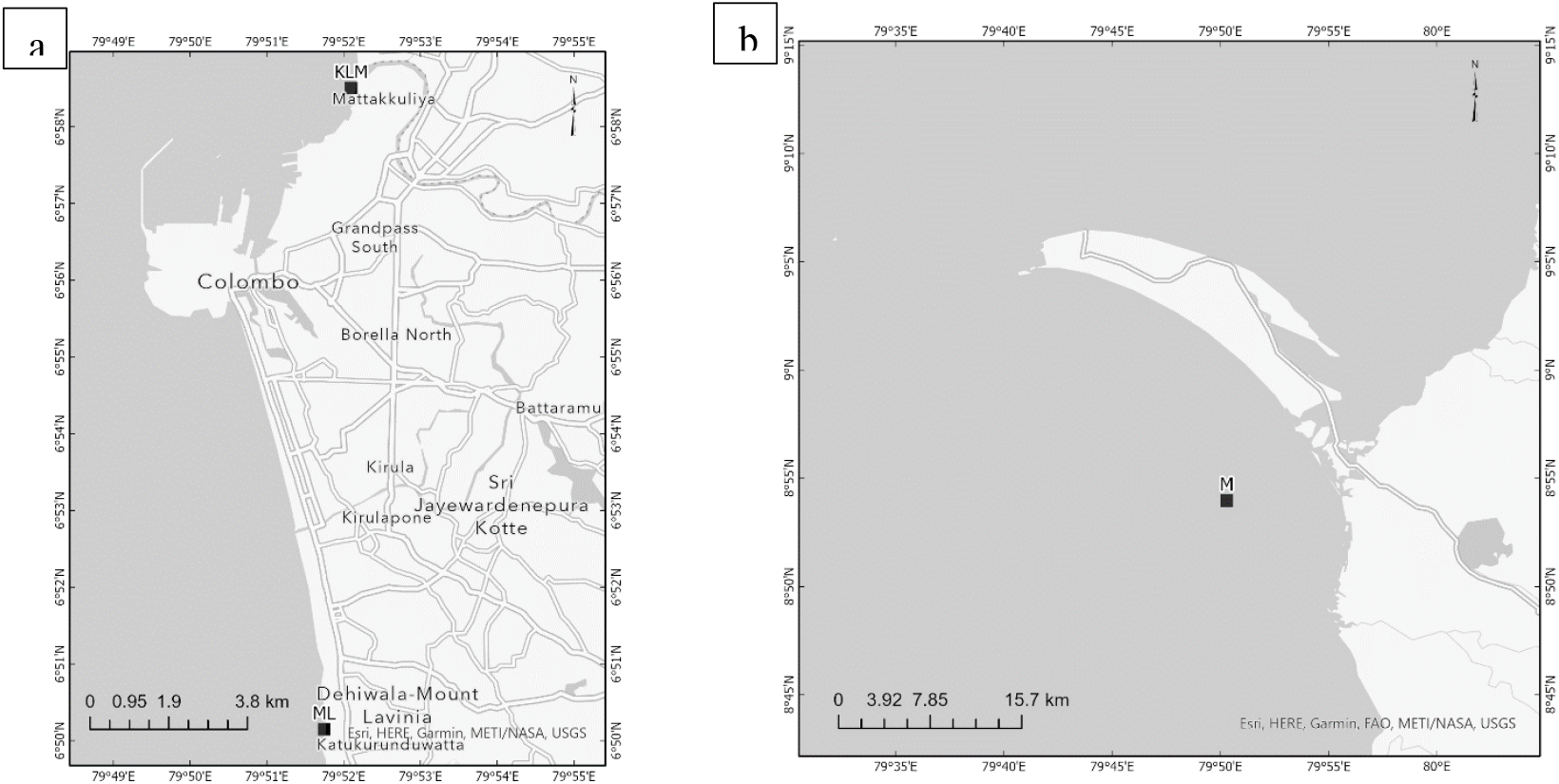
Reference sampling locations (a) coastal waters outside the Colombo Port (b) Mannar.

The sampling in the port was carried out using settlement structures initially. Following the preliminary field surveys, three types of settlement collectors were designed with different substrates attached to enhance the settlement of fouling organisms. The first two types (i and ii) of structures were made of PVC pipes connected by four elbows at the corners and one tee joint to pass the rope. Plates (10×10 cm and 10×20 cm) made of steel sheets coated with an Al-Zn alloy polyester-based coating were attached to them (Figure 2.3). The third settlement structure (iii) had four types of panels made of canvas (a), non-coated iron net (b), coir ropes (c) and epoxy paint coated iron net (d), respectively, and was deployed in the selected sampling stations. Settlement collectors were retrieved at monthly intervals for two years and recorded with digital close-up photographs for further analysis. Reference fouling samples were collected from ML and KLM using 20 m2 belt transects. Similarly, dive sampling was conducted up to a 5 m depth in the coastal waters off Mannar to collect sponges. The samples were stored in sealed bags and transported to the laboratory under chilled conditions for identification and further analysis.

**Figure: 2.3.**
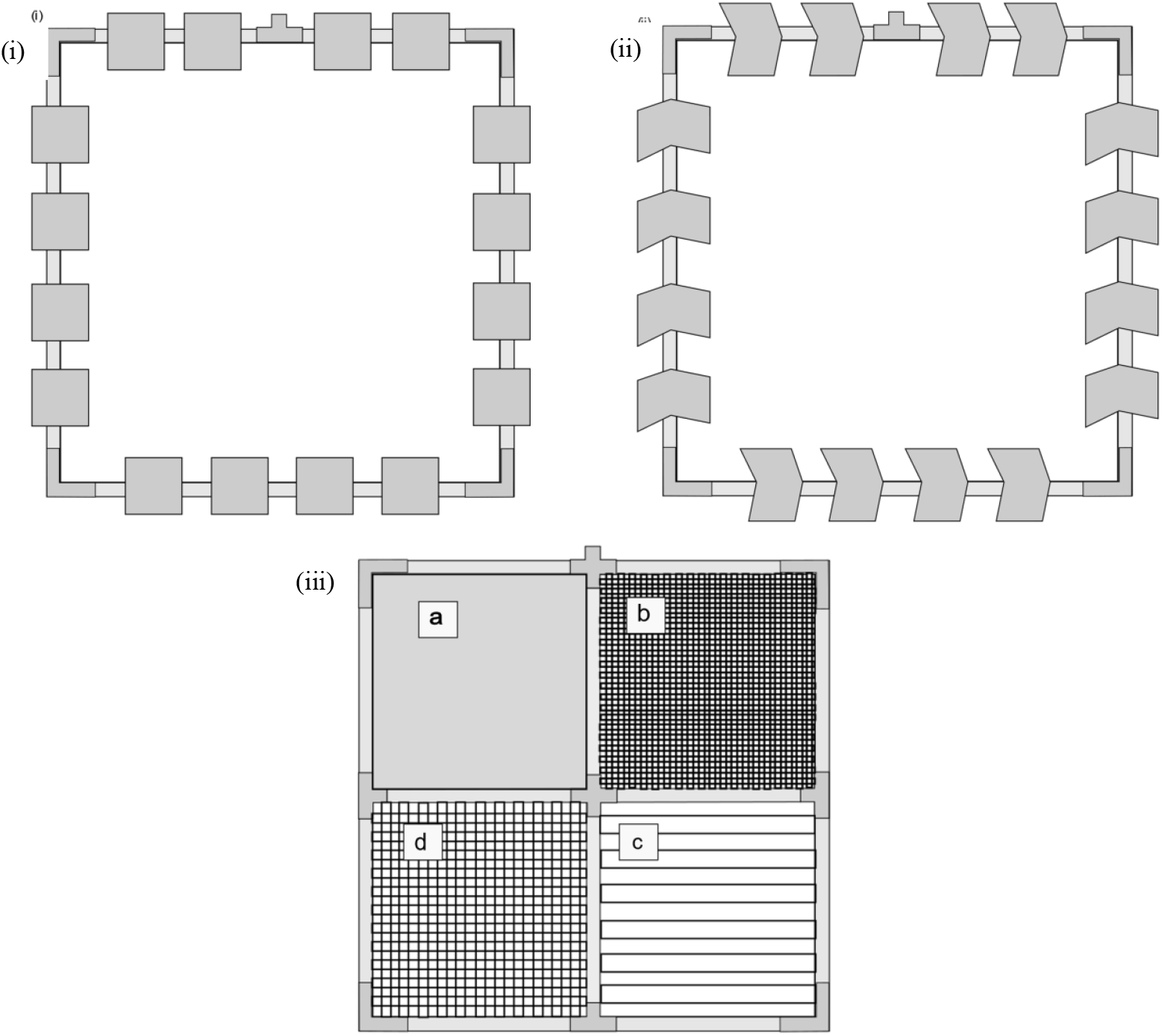
Settlement collectors made up of PVC frames fitted with metal plates. (i) with 10×10 cm plates and (ii) with 10×20 cm plates. (iii) and a settlement structure with four different substrates.

### 2.1. Sample selection and extract preparation

All the species identities were confirmed with the help of species identification databases and taxonomic literature (*General Shell Portal*, 2020; Marasinghe, 2019; *WoRMS - World Register of Marine Species*, 2019). Furthermore, sponge species were identified to the lowest taxonomic level by studying their spicule patterns under a zoom stereo microscope (Nikon SMZ 1270i). In addition, SEM images of sponge spicules were obtained when further confirmation was needed (S1). *Schizoporella errata, Botrylloides violaceus, Carijoa riisei, Callyspongia diffusa*, and *Mycale parishii* were selected from the port (S2) for the experiment after an extensive screening process. Colonies of bryozoan *S. errata* were collected from all the sampling locations, while sponges *C. diffusa* and *M. parishii* were extracted from PJ and BQ. *C. riisei* soft coral was collected from UCT and PJ, and tunicate *B. violaceus* was collected from PJ, UCT, and CICT. Specimens from each species were preserved for future reference. Simultaneously, *Halichondria panicea* from the KLM and *Hymeniacidon perlevis* from ML were selected, along with *Acanthella cavernosa* and *Clathria* sp. (S3) from the sampling location at Vankalei, Mannar.

The selected organisms were rinsed with fresh water to remove debris and lyophilized at the laboratory. Afterwards, weighed samples were ground and soaked in 25 mL of HPLC grade methanol. Both maceration and sonication at room temperature were used to extract both inter- and intracellular metabolites from the samples. The solvent level was maintained higher than the sample level at all times and the process was continued for 3 days with 24 h extraction cycles (Annegowda et al., 2012; Ebada et al., 2008). Excess methanol was evaporated in the rotary evaporator. The dry product was processed following the experimental procedures mentioned in the literature for the anti-settlement assay followed by phytochemical screening and GC-MS analysis (Muzquiz, 2000; Peters et al., 2004; Wesołowska, et al., 2016).

### 2.2. Anti-settlement assay

Following the findings by Mannapperuma et al. (2013), *Escherichia coli* bacterial strains were selected as a representation of the microbial settlement in the port environment. The assay was conducted using the method described in Larimer et al. (2015) with minor modifications. First, a broth culture of *E. coli* (ATCC 25922) with an OD_600_ = 0.35 was prepared for the experiment. Then the coating was arranged using previously prepared dried chemical extractions, dimethyl sulfoxide, and epoxy paint. The final concentrations of each coating were set at 100 μg mL^−1^. Coupons (2.5 cm × 2.5 cm) made of steel plates were prepared to examine the effect. Each coupon was covered with the respective coatings prepared using the test species and placed in a petri dish containing 25 mL of LB broth. Each dish contained two coupons: one for cell density measurements and the other for further experiments. Finally, an inoculum of 100 μL of *E. coli* was introduced and kept covered at 29 °C in a stationary position.

A single coupon from each concentration was removed and analysed at six intervals for three days (0, 12, 24, 36, 48, and 60 h). The experiment was triplicated for each time point. Removed coupons were dipped in a phosphate-buffered saline (PBS) solution for 1-2 s in order to remove unbound fouling bacteria and organic matter. Following that, samples at each time point were placed in 50 mL falcon tubes containing 15 mL of PBS with 0.05% Triton X-100. Bacteria attached to the coupons were dispersed with sonication for 30 min, followed by vortexing before the analysis. Three 1 mL samples were pipetted out to measure optical density at 600 nm (OD_600_) using UV-visible spectroscopy (JENWAY 6305) for each sample. A basic assumption was made that the settlement of *E. coli* is directly proportional to the optical density, and the results were analysed to determine the anti-settlement property of individual extractions using Two-way ANOVA.

### 2.3. Preliminary phytochemical screening and GC-MS analysis

#### 2.3.1. Preliminary phytochemical screening

A fraction of the dry extract was dissolved in ethanol to obtain 200 mL of a 1 mgmL^−1^ solution. Methods described in Yusuf et al. (2014) and Bargah (2015) were used to qualitatively determine the presence of alkaloids, sterols, triterpenes, saponins, phenols, and flavonoids during the process.

#### 2.3.2. GC-MS analysis

Volatile and semi-volatile component analysis was conducted to find out the presence of secondary metabolites composition subsequently to preliminary phytochemical screening. The species were extracted following the methods mentioned by Ebada et al. (2008) with some modifications to produce crude hexane, dichloromethane (DCM) and methanol extracts, with DCM replacing ethyl acetate during the extraction. The extracts were then dried and combined. The final dry product was divided into three parts, one of which was redissolved in dichloromethane (2 mL) for GC-MS analysis. Chromatographic analysis was repeated on the dry products of three solvent fractions separately when noticeable representations were observed.

Volatile component analysis was conducted on an Agilent 7890A GC coupled with an Agilent 5975C quadrupole MS with Triple-Axis Detector. Dichloromethane extracts of every sponge species were analysed following a similar method. The prepared extract was subjected to centrifugation at 10000 rpm for 30 minutes. The supernatant was filtered through Whatman 42.5 micron GF/C filter paper. The flow rate of the carrier gas (He) was maintained at 1 mL min^−1^ and 4 μL of the sample was injected at a split ratio of 5: 1. The injector temperature and the transfer line were set at 280 °C and the source temperature was kept at 230 °C. The column was kept for 5 min at the initial temperature of 40 °C then increased to 60 °C at a rate of 6 °C min^−1^. After 10 min, the temperature was increased to 280 °C at a rate of 30 °C min^−1^. The temperature was maintained at 280 °C for 40 min. The mass spectra were recorded at a standard source electron energy of 70 eV (Wesołowska, et al., 2016). Subsequently, dichloromethane extracts of soft coral *Carijoa riisei* and tunicate *Botrylloides violaceus* were filtered and prepared for the analysis following the method described previously, and Helium (99.999%) was used as carrier gas at a constant flow rate of ± 1 mL mn^−1^. An injector volume of 2 μL with a split ratio of 10:1 was injected, and the injector temperature and source temperature were set at 250 °C and 280 °C respectively. The oven temperature was programmed from 110 °C to 200 °C for 2 min at a 10 °C min^−1^ rate, then increased to 280 °C at 5 °C min^−1^ and maintained for 9 min (Meenakshi et al., 2013).

Next, the crude extract of *S. errata* was centrifuged for 30 min at 10000 rpm and filtered through Whatman 42.5 micron GF/C filter paper. The temperature programme was carried out according to Muzquiz (2000) and Peters et al. (2004) with modifications. 1 μL of *S. errata* dissolved in dichloromethane was injected into the GC-MS at a 5:1 split ratio. The carrier gas (He=99.999%) flow rate was 1 mL min^−1^. The oven temperature was increased from 90 °C to 160 °C at 6 °C min^−1^ and maintained for 10 min. After that, the temperature was increased to 280 °C at a 10 °C min^−1^ rate and maintained for 30 min. The injector temperature was set at 240 °C.

## 3. RESULTS

### 3.1. Anti-settlement Assay

Optical density values of the anti-settlement assay for target species, including the control, stabilised after the 48h time point. (Figure 3.1). The absorption values of all the species portrayed against time showed fairly similar behaviour towards *E. coli* settlement. Individual optical density (OD) values of each organism were visibly different from each other and were lower in comparison to the control. OD values for the first 12h for *S. errata, B. violaceus*, and *C. diffusa* were below 0.1 compared to the rest of the species harvested from Colombo port.

Similarly, for two sponge species collected from adjacent coastal waters as well as sponge A. cavernosa collected from Mannar, OD values recorded in the first 12 hours were below 0.1. Analysis of absorbance data using Minitab, Two-way ANOVA indicated the anti-settlement property of individual extractions is not significantly different (p = 0.235 > 0.05). However, there is a significant difference in the anti-settlement activity between target organisms and control in terms of fraction types (p = 0.000 < 0.05) and experimental duration (p = 0.047 < 0.05).

**Figure 3.1.**
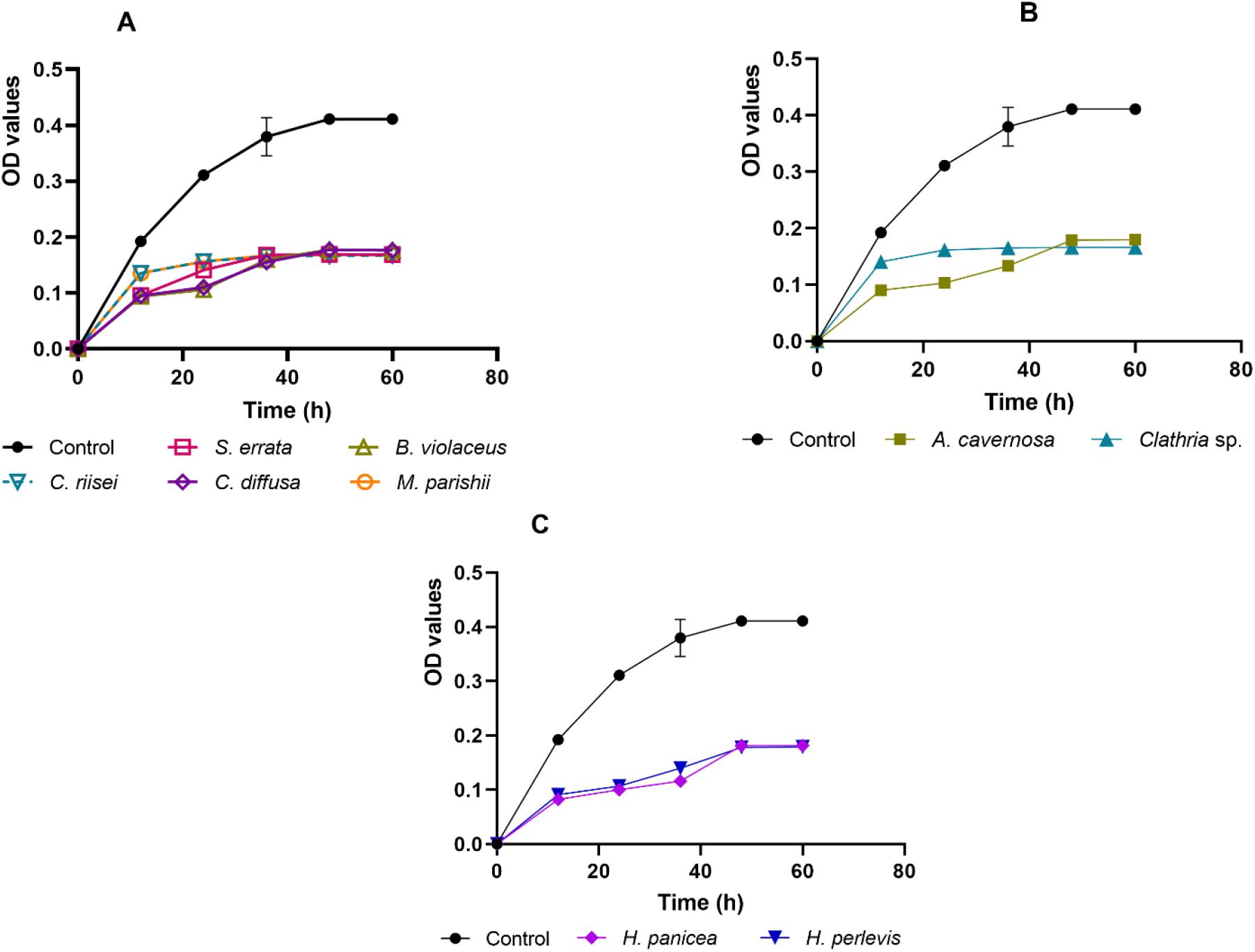
OD values of target organisms for the anti-settlement assay.

A-species collected from Colombo Port, B-species from Mannar and C-species from coastal waters outside the Colombo Port.

#### 3.2. Preliminary phytochemical screening and GC-MS analysis

### 3.2.1. Preliminary phytochemical screening

Steroids and terpenoids were identified as the two most common metabolites found in test organisms. *Carijoa riisei, Acanthella cavernosa, Callyspongia diffusa, Halichondria panicea*, and *Hymeniacidon perlevis* indicated a strong presence of steroids and terpenoids in a dark reddish-brown colour. Similarly, saponins were detected for all the species, with *C. Diffusa* showing a noticeable reaction. While alkaloids were found to be a common metabolite in the majority of the organisms studied, they were noticeably absent in *Botrylloides violaceus*. None of the species have been observed with phenols, tannins or flavonoids. Among all the species, *C. diffusa* has given the most noteworthy reaction, while *B. violaceus* has shown the weakest response of all the species to the phytochemical screening (Table 3.1).

**Table 3.1.**
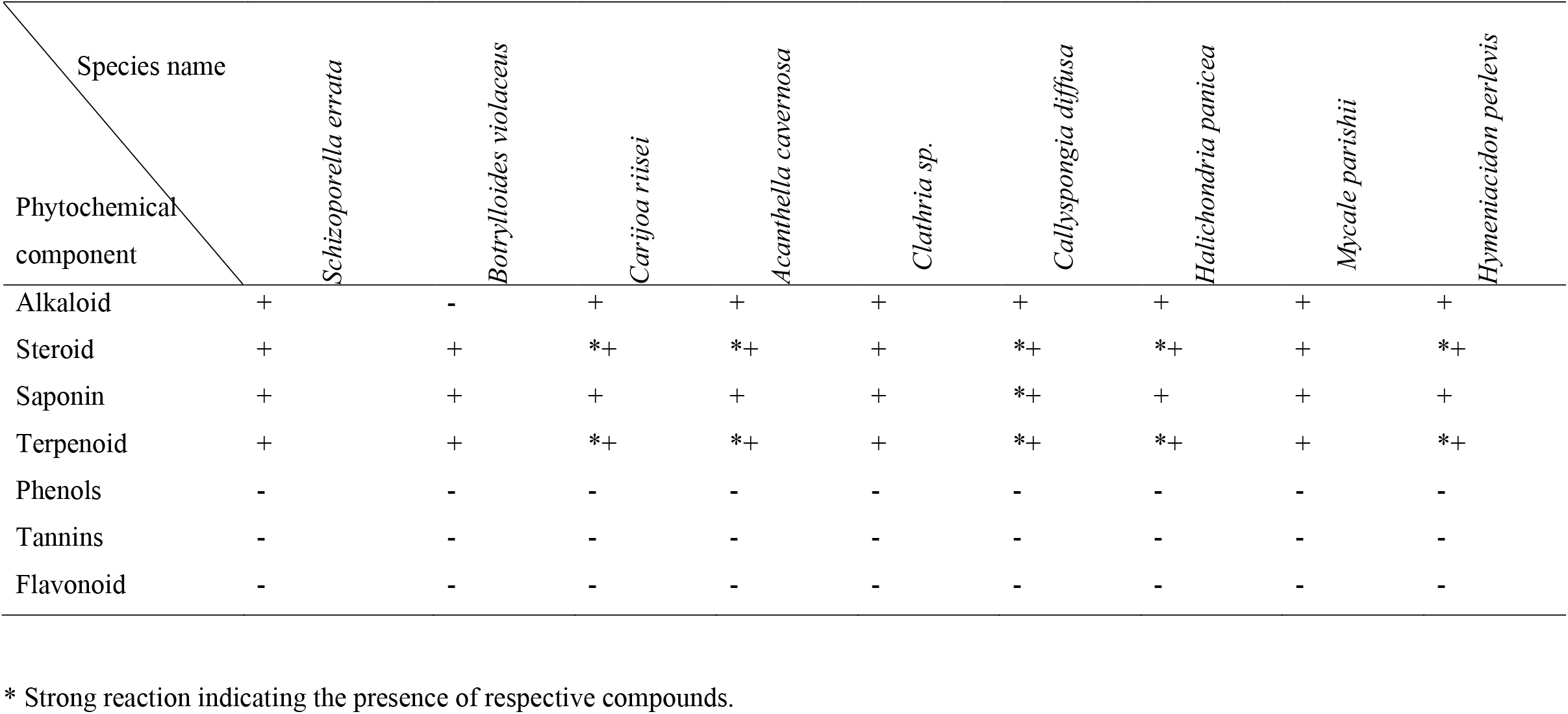
Phytochemical presence in methanolic extracts of *Schizoporella errata, Botrylloides violaceus, Carijoa riisei, Acanthella cavernosa, Clathria* sp., *Callyspongia diffusa, Halichondria panicea, Mycale parishii* and *Hymeniacidon perlevis*.

| Species name<br>Phytochemical<br>component | *Schizoporella errata* | *Botrylloides violaceus* | *Carijoa riisei* | *Acanthella cavernosa* | *Clathria* sp. | *Callyspongia diffusa* | *Halichondria panicea* | *Mycale parishii* | *Hymeniacidon perlevis* |
| --- | --- | --- | --- | --- | --- | --- | --- | --- | --- |
| Alkaloid | + | - | + | + | + | + | + | + | + |
| Steroid | + | + | *+ | *+ | + | *+ | *+ | + | *+ |
| Saponin | + | + | + | + | + | *+ | + | + | + |
| Terpenoid | + | + | *+ | *+ | + | *+ | *+ | + | *+ |
| Phenols | - | - | - | - | - | - | - | - | - |
| Tannins | - | - | - | - | - | - | - | - | - |
| Flavonoid | - | - | - | - | - | - | - | - | - |

\* Strong reaction indicating the presence of respective compounds.

### 3.2.2. GC-MS analysis

GC-MS analysis for the 9 target organisms followed by phytochemical screening has characterised the occurrence of secondary metabolites. In GC-MS spectra, alkaloid compounds that were present in preliminary phytochemical screening were absent, while the phenolic compounds that were previously absent were detected. Noticeably, compounds from *Hymeniacidon perlevis* were indefinable due to the noise that was present in the GC-MS spectrum. The majority of the resulting compounds were esters, followed by fatty acids, sterols, alkanes and terpenes. Some of the organisms showed similarities in chemical profiles by which *C. diffusa* and *S. errata* exhibited the highest number of esters detected, while *S. errata* spectrum was observed with the presence of a significant number of fatty acid compounds. The majority of sterol compounds were obtained from *Clathria* sp., followed by *C. diffusa* with a high number of terpenoid and phenolic compounds. Among the experimented individuals, *C. diffusa* was accentuated for having a great variety of secondary metabolites. 13 known compounds with a higher degree of antibacterial activity were identified among the collective of secondary metabolites.

Octadecanoic acid is a saturated long-chained fatty acid observed in *S. errata* and *C. diffusa* chromatograms. It is commonly known as stearic acid. Similarly, tetradecanoic acid, which is known as myristic acid, was observed in both *S. errata* and *C. diffusa*. Both species further exhibited the presence of dodecanoic acid in their chromatograms for the GC-MS analysis. In addition to these two species, dodecanoic acid was observed in the sponge *M. parishii* and the marine tunicate *B. violaceus*. Furthermore, except for tunicate *B. violaceus*, the other three species have given a positive indication of hexadecenoic acid presence. Apart from that, *C. riisei* and *A. cavernosa* were also observed with hexadecenoic acid.

Eicosane is a straight-chain alkane observed in volatile component analysis of sponges, *Clathria* sp., and *M. parishii*. A few sterols were present in the GC-MS analysis of the target species. Among the collective of sterols, γ-sitosterol (clionasterol), a compound with 29 carbon atoms, was detected in the spectrum of *Clathria* sp. Likewise, the presence of a widely known 3beta-sterol, Ergosta-5,22-dien-3-ol, (3β,22E) (brassicasterol) was also observed in *Clathria* sp. A sesquiterpene compound, caryophyllene, was observed in both *C. diffusa* and *M. parishii* sponges. Among the test species, *C. diffusa* can be recognised as a species with a number of significant secondary metabolites. In the chromatogram of *C. diffusa*, several important compounds were further observed. It was observed with two non-cyclic monoterpenoids, 1,6-Octadien-3-ol, 3,7-dimethyl (linalool) and 2,6-Octadien-1-ol, 3,7-dimethyl-, (E)-(geraniol), as well as a widely known bioactive sesquiterpene compound 1S-1α,2β,4β-1-Ethenyl-1-methyl-2,4-bis-(1-methylethenyl)cyclohexane (β-elemene), and a known bioactive phenylpropanoid compound called (Z)-5-Propenyl-1,2,4-trimethoxybenzene (β-asarone). Furthermore, the sponge *H. panicea* was observed with the rare long-chain alkane hydrocarbon hentriacontane.

## 4. DISCUSSION

An anti-settlement assay was conducted in order to inquire about the effect of each extract on the settlement of *Escherichia coli*. The direct effect of each extract can be interpreted following the basic assumption made at the beginning, that *E. coli* settlement is directly proportional to the optical density. The manner of *E. coli* settlements on the coupons was observed similarly for *Carijoa riisei, Mycale parishii*, and *Clathria* sp. Among the three, *C. riisei* and *M. parishii* were collected from Colombo Port, where the human influence is high, and *Clathria* sp. was collected from the reference point with the least influenced environment; Mannar. The settlement inhibition values of *Callyspongia diffusa, Acanthella cavernosa, Halichondria panicea*, and *Hymeniacidon perlevis* were also similar to each other. Two species: *H. panicea* and *H. perlevis*, were collected from reference sampling points in coastal waters closer to Colombo Port. *C. diffusa* was collected from the PJ of Colombo Port; and sponge *A. cavernosa* was collected from Mannar.

Individual OD values closely resembled each other except for the three species: *H. parishii, S. errata, and C. diffusa*, which were collected from Colombo Port. Furthermore, the biofilm formation of *E. coli* was interrupted by *B. violaceus* and *S. errata* similarly at the beginning (OD below 0.1). Likewise, *S. errata, B. violaceus, C. diffusa, A. cavernosa, H. panicea*, and *H. perlevis* showed a low rate of *E. coli* settlement in the first 12 h in comparison to that of *C. riisei, M. parishii*, and *Clathria* sp. However, during the second 12 h period, *E. coli* settlement was relatively similar for all the species except the tunicate *B. violaceus* and bryozoan *S. errata*. The experiment results indicate that the affective nature of the extracts did not vary with harvesting location or environment.

*B. violaceus* is known inhibitor of the larval recruitment, which explains the noticeable anti-settlement activity of the current study (Dijkstra et al., 2007). *H. panicea’*s individual anti-settlement appears similar to *S. errata* and *Clathria* sp. However, in comparison of OD values during the first 12 h of the experiment between *H. panicea* (OD < 0.1), *Clathria* sp. (OD > 0.1) and *S. errata* (OD < 0.1), it was shown that *S. errata* crude extraction efficacy is coequal to *H. panicea* on *E. coli* settlement at the beginning. Most importantly, three out of the five species collected from the port, namely, *S. errata, C. riisei*, and *B. violaceus*, are considered globally known invasive species according to the Global Invasive Species Database (GISD). However, all three of them showed corresponding anti-settlement activity without noticeable deviations from each other.

Secondary metabolite analysis has resulted in a collective of chemical compounds that possibly explains the anti-settlement assay. They are important chemical compound producers in the world, playing a great role in possessing antibacterial, anticancer, anti-infective, and anti-inflammatory properties. The noticeable reaction of preventing biofilm formation indicates that the respective chemical groups are present in greater quantities in the species studied. The chemical richness of sponges appears to be the same for all the species, despite being collected from different locations. Triterpenoids, aromatic acids, and steroids were reported previously in a phytochemical screening conducted on *C. diffusa* and *M. parishii (Zygomycale parishii)* collected from the southwest coast of India (Krishnan & Keerthi, 2016). In addition to that, the presence of phenolic compounds was also reported for these two species. However, phenolic compound presence was not detected for any of the species, including *C. diffusa* and *M. parishii*, for the present study in phytochemical screening. Similarly, *S. errata, B. violaceous*, and *C. riisei* indicated the presence of the above-mentioned chemical groups.

The 13 chemical compounds identified in the study were previously identified to have properties that potentially reduce or terminate biofilm formation. They have been shown to have antibacterial, anti-cancer, anti-inflammatory, antifungal, and larvicidal properties against other organisms (Abubakar & Majinda, 2016; Bai A & Vittal, 2014). Octadecanoic acid shown in the marine sponge *C. diffusa* and the marine bryozoan *S. errata* is a long-chained fatty acid, which is often described as an intermediate for a wide range of commercial purposes such as resins, stabilizers, and lubricants. The presence of octadecanoic acid in a marine organism was previously reported by Si et al. (2001) in the sponge *Callyspongia fibrosa*. The compound has been reported to have antimicrobial activity by Abubakar and Majinda (2016). Both *S. errata* and *C. diffusa* further exhibited the presence of fatty acids and tetradecanoic acid (myristic acid) in their chromatograms. According to Chen et al. (2019), myristic acid appeared to influence cell death and inhibited growth when its effect was examined against *Listeria monocytogenes* in milk.

Dodecanoic acid (lauric acid) is known to exhibit antibacterial properties. Huang et al. (2014) further discussed the anti-inflammatory property of dodecanoic acid against the bacteria *Propionibacterium acnes*, which is a possible cause of acne inflammation. In addition to that, Kwan et al. (2011) explained the quorum sensing inhibition of dodecanoic acid, which was reported to be of a higher and similar magnitude to the known quorum sensing inhibitor malyngolide. The chemical profile of *S. errata* exhibited the presence of myristic acid and lauric acid, which fortifies the bioactive nature of *S. errata* against *E. coli* settlement. *S. errata* anti-settlement activity is equivalent to that of *C. diffusa* within the first 12 hours of the experiment and noticeably maintained at a low rate until the end, which signifies the complementary effect of bioactive compounds in its possession. In addition to that, both *B. violaceus* and *M. parishii* exhibited the presence of dodecanoic acid. The presence of methyl laurate in the marine microalga *Oscillatoria* sp. was previously described by Son et al. (2000). According to the (European Chemical Agency, 2020) methyl laurate is very toxic to aquatic life with long-lasting effects in which quorum sensing inhibition to a certain degree was displayed in different marine cyanobacteria. Thus, it can reinforce the anti-settlement property of *B. violaceus*, although the tunicate appears to have low chemical diversity compared to the sponge chemical compositions.

Shaaban et al. (2021) describe the high inhibitory activity of clove extracts containing methyl palmitate (hexadecanoic acid methyl ester) against multidrug-resistant bacteria. Furthermore, the anti-inflammatory property was expressively described by Saeed et al. (2012) in their study. This particular chemical compound presence was detected in five species that were considered for the volatile compound analysis, except *B. violaceus, H. panicea*, and *Clathria* sp. During the first 12 h of the assay, sponges *A. cavernosa, C. diffusa*, and *M. parishii*, which contained methyl palmitate, showed comparatively high anti-settlement activity compared to *Clathria* sp. The fatty acids were converted prior to the GC-MS analysis. Therefore, the possibility of the involvement of palmitic acid (hexadecanoic acid)-like compounds in the inhibition of biofilm formation can be further strengthened.

Krishnan and Keerthi (2016) mentioned the secondary metabolite analysis of *M. parishii* collected from the southwest coast of India. According to them, eicosane was reported in methanol and dichloromethane extracts of *M. parishii*. Eicosane (icosane) was also recorded in both *Clathria* sp. and *M. parishii* in the current study for the volatile component analysis. The above two species were harvested from two separate locations, yet the eicosane presence can complement the environmental resilience with its antifungal property. Furthermore, the chromatogram of *Clathria* sp. has resulted in the presence of a number of sterols in contrast to the minimal presence of fatty acids. *Clathria* sp. spectrum has exhibited the presence of clionasterol, a plant and marine metabolite. Some of the compounds, like clionasterol, have been previously reported in other marine organisms. Santalova et al. (2004) mentioned the existence of clionasterol as the main sterol of the sponge *Clathria major*. Similarly, brassicasterol, which was seen in the *Clathria* sp. chromatogram, has been described previously with antimicrobial properties. Hassan (2020) describes the anti-infective property of brassicasterol against herpes simplex virus type 1 (HSV-1) and *Mycobacterium tuberculosis* (Mtb).

Caryophyllene (β-caryophyllene) is a natural bicyclic sesquiterpene. It is one of the significant findings obtained for both *C. diffusa* and *M. parishii* in the current study. The compound is commonly found in aromatic oils like clove and rosemary oils. The compound is also described as a non-steroidal anti-inflammatory agent (National Center for Biotechnology Information, 2019). Marine-derived caryophyllene-related compounds were previously recorded in soft coral *Sinularia nanolobata*, gorgonian coral *Subergorgia suberosa, the* marine fungus *Humicola fuscoatra*, and the sponge-associated fungus *Hansfordia sinuosa* (Wu et al., 2014). They were further characterised as having weak cytotoxic and inhibitory effects against four bacteria, including *Escherichia coli*. In addition, the strong local anaesthetic ability of β-caryophyllene was also reported by Ghelardini et al. (2001), and it was further discussed as a possible quorum sensing inhibitor in the study of Bai A and Vittal (2014).

GC-MS analysis has also indicated the occurrence of terpenoids in the target individuals. One of the known plant metabolites, linalool, was observed in the sponge *C. diffusa* extracts. Herman et al. (2016) report the use of linalool to maintain the microbial purity of products such as cosmetics and medicine due to its ability to enhance the antimicrobial effectiveness of products. Furthermore, the *C. diffusa* chromatogram has also exhibited the presence of a monoterpenoid called geraniol and a sesquiterpene, β-elemene. Holst et al. (1994) and Ichikawa et al. (1993) mentioned the existence of geraniol as well as its derivatives in *Halichondria* sp. and the bryozoan *Flustra foliacea*. Nevertheless, volatile component analysis of *H. panicea* has not resulted in geraniol or its derivatives in the current study. The effectiveness of the compounds was further discussed by Takamura et al. (2013). According to them, hybrid molecules consisting of geraniol and butenolide enhanced the antifouling activity. In addition, the remarkable antimicrobial activity of β-elemene was discussed against *Mycobacterium tuberculosis* by Sieniawska et al. (2018). The occurrence of bioactive products in *C. diffusa* extracts corresponds to its impact on the biofilm formation of *E. coli. C. diffusa* exhibited significant inhibition against the bacterial settlement within the first 24 h compared to other individuals in which its performance was only positioned behind *H. perlevis, H. panicea*, and *A. cavernosa*.

β-asarone shown in the *C. diffusa* chromatogram is one of the pharmacologically noteworthy chemical compounds. The antimicrobial property of asarone was previously mentioned in the study of Mukherjee et al. (2007). In addition, a few other bioactive properties such as anti-depressant, anticonvulsant, anti-inflammatory, antioxidant, and hypolipidemic were discussed in detail in several studies (Li & Wah, 2017; Mukherjee et al., 2007; Uebel et al., 2020). Phenylpropanoid compounds, as well as known potent antimicrobial agents such as sesquiterpenes, naturally enhance their bioactive properties (Sieniawska et al., 2018). Thus, strengthening the possible reaction of C. diffusa against biofilm formation of E. coli. Moreover, the sponge *H. panicea* has exhibited the presence of an alkane called hentriacontane for the volatile component analysis. Hentriacontane is known for its anti-microbial, anti-inflammatory, and anti-tumor properties (Khajuria et al., 2017; Takahashi et al., 1995). The difference between each extract and the settlement of *E. coli* was not significant. However, the comparison of bacterial-settlement at each time point versus individual extractions appears to be slightly different. *H. panicea* has expressed the most noteworthy response within the first 24 h of the experiment out of all the species. Soft coral, *C. riisei*, as well as two sponges, *M. parishii* and *Clathria* sp., have shown a high degree of *E. coli* settlement compared to others. In comparison to the less chemically rich *C. riisei* and *M. parishii*, the inhibition of biofilm formation of the sponge *Clathria* sp. contained a number of secondary metabolites that appeared weaker in the experiment. Likewise, *B. violaceus* was observed with a few noteworthy secondary metabolites in possession, nevertheless able to demonstrate a remarkable anti-settlement property during the current experimental conditions. The destruction of chemical compounds during the extraction procedure as well as the cloaking effect caused by chemically or physically similar compounds might have produced such results. The overall effect of each extract confirms the anti-settlement activity of each individual against *E. coli* bacteria, hence providing strong evidence of the biofilm formation inhibition properties of the species considered. However, the chemical nature of the existing compounds suggests the combined effect on the inhibition of biofilm formation. Therefore, further research is needed to assess the precise role of each chemical compound on biofilm formation.

## Acknowledgements

This work was funded by University Research Grant ASP/01/RE/SCI/2015/28. The experiments were facilitated by the Department of Zoology, Department of Chemistry and Instrument Center of the Faculty of Applied Sciences (IC-FAS) of the University of Sri Jayewardenepura (USJP). The authors gratefully acknowledge, G. C. Madushanka of Faculty of Graduate Studies for providing necessary maps, K. Vitharana and P. Suganda of Asian Paints Lanka Ltd for providing epoxy coatings.

## LITERATURE CITED

Abubakar, M. N., & Majinda, R. R. T. (2016). GC-MS analysis and preliminary antimicrobial activity of Albizia adianthifolia (Schumach) and Pterocarpus angolensis (DC). Medicines (Basel, Switzerland), 3(1), 3. https://doi.org/10.3390/medicines3010003

Annegowda, H. V., Bhat, R., Min-Tze, L., Karim, A. A., & Mansor, S. M. (2012). Influence of sonication treatments and extraction solvents on the phenolics and antioxidants in star fruits. Journal of Food Science and Technology, 49(4), 510–514. https://doi.org/10.1007/s13197-011-0435-8

Bai, A. J., & Vittal, R. R. (2014). Quorum sensing inhibitory and anti-biofilm activity of essential oils and their in vivo efficacy in food systems. Food Biotechnology, 28(3), 269–292. https://doi.org/10.1080/08905436.2014.932287

Bargah, R. (2015). Preliminary test of phytochemical screening of crude ethanolic and aqueous extract of Moringa pterygosperma Gaertn. Journal of Pharmacognosy And Phytochemistry, 4(1):07–09.

Bekiari, V., Nikolaou, K., Koromilas, N., Lainioti, G., Avramidis, P., Hotos, G., Kallitsis, J. K., & Bokias, G. (2015). Release of Polymeric Biocides from Synthetic Matrices for Marine Biofouling Applications. Agriculture and Agricultural Science Procedia, 4, 445–450. https://doi.org/10.1016/j.aaspro.2015.03.051

Callow, M. E., & Callow, J. E. (2002). Marine biofouling: A sticky problem. Biologist (London, England), 49(1), 10–14.

Chen, X., Zhao, X., Deng, Y., Bu, X., Ye, H., & Guo, N. (2019). Antimicrobial potential of myristic acid against Listeria monocytogenes in milk. The Journal of Antibiotics, 72(5), 298–305. https://doi.org/10.1038/s41429-019-0152-5

Clarke Murray, C., Therriault, T. W., & Pakhomov, E. (2013). What lies beneath? An evaluation of rapid assessment tools for management of hull fouling. Environmental Management, 52(2), 374–384. https://doi.org/10.1007/s00267-013-0085-x

Devendra, S., & Muthucumarana, R. (2013). Maritime Archaeology and Sri Lanka: Globalization, immigration, and transformation in the underwater archaeological record. Historical Archaeology, 47(1), 50–65. https://doi.org/10.1007/BF03376889

Dijkstra, J., Sherman, H., & Harris, L. G. (2007). The role of colonial ascidians in altering biodiversity in marine fouling communities. Journal of Experimental Marine Biology and Ecology, 342(1), 169–171. https://doi.org/10.1016/j.jembe.2006.10.035

Drake, J. M., & Lodge, D. M. (2007). Hull fouling is a risk factor for intercontinental species exchange in aquatic ecosystems. Aquatic Invasions, 2(2), 121–131. https://doi.org/10.3391/ai.2007.2.2.7

Ebada, S. S., Edrada, R. A., Lin, W., & Proksch, P. (2008). Methods for isolation, purification and structural elucidation of bioactive secondary metabolites from marine invertebrates. Nature Protocols, 3(12), 1820–1831. https://doi.org/10.1038/nprot.2008.182 PMID:18989260

European Chemical Agency. (2020). Substance Information-Methyl laurate. ECHA. https://echa.europa.eu/substance-information/-/substanceinfo/100.003.556

Ferreira-Vançato, Y. C., Dantas, F. M., & Fleury, B. G. (2020). Nanobiocides against marine biofouling. Studies in Natural Products Chemistry, 67, 463–514. https://doi.org/10.1016/B978-0-12-819483-6.00013-8

Floerl, O., Inglis, G. J., Dey, K., & Smith, A. (2009). The importance of transport hubs in stepping-stone invasions. Journal of Applied Ecology, 46(1), 37–45. https://doi.org/10.1111/j.1365-2664.2008.01540.x

General Shell Portal. (2020). Idscaro. http://www.idscaro.net/sci/

Ghelardini, C., Galeotti, N., Mannelli, L. D. C., Mazzanti, G., & Bartolini, A. (2001). Local anaesthetic activity of β-caryophyllene. Il Farmaco, 56(5-7), 387–389.

Hassan, S. T. (2020). Brassicasterol with dual anti-infective properties against HSV-1 and Mycobacterium tuberculosis, and cardiovascular protective effect: nonclinical in vitro and in silico assessments. Biomedicines, 8(5), 132.

Herman, A., Tambor, K., & Herman, A. (2016). Linalool Affects the Antimicrobial Efficacy of Essential Oils. Current Microbiology, 72(2), 165–172. https://doi.org/10.1007/s00284-015-0933-4

Hewitt, C. L., Everett, R. A., & Parker, N. (2009). Examples of current international, regional and national regulatory frameworks for preventing and managing marine bioinvasions. In Biological invasions in marine ecosystems (pp. 335–352). Springer, Berlin, Heidelberg.

Holst, P. B., Anthoni, U., Christophersen, C., Nielsen, P. H., & Bock, K. (1994). A racemic diterpene from the marine bryozoan Flustra foliacea, natural product or artefact? Acta Chemica Scandinavica, 48(9), 765–768.

Huang, W. C., Tsai, T. H., Chuang, L. T., Li, Y. Y., Zouboulis, C. C., & Tsai, P. J. (2014). Anti-bacterial and anti-inflammatory properties of capric acid against Propionibacterium acnes: a comparative study with lauric acid. Journal of dermatological science, 73(3), 232–240.

Ichikawa, Y., Yamazaki, M., & Isobe, M. (1993). Novel, regioselective allylamine construction; first synthesis of geranyllinaloisocyanide, a diterpene from the marine sponge, Halichondria sp. Journal of the Chemical Society, Perkin Transactions 1, (20), 2429–2432.

Jayasundara, R. M., & Ranatunga, R. R. M. K. P. (2016). Macro-Fouling Faunal Assemblage in Hambantota Port. In Proceedings of International Forestry and Environment Symposium (Vol. 21).

Khajuria, V., Gupta, S., Sharma, N., Kumar, A., Lone, N. A., Khullar, M., Dutt, P., Sharma, P. R., Bhagat, A., & Ahmed, Z. (2017). Anti-inflammatory potential of hentriacontane in LPS stimulated RAW 264.7 cells and mice model. Biomedicine & Pharmacotherapy, 92, 175–186. https://doi.org/10.1016/j.biopha.2017.05.063

Kim, A., Kim, J. H., & Patel, R. (2021). Modification strategies of membranes with enhanced Anti-biofouling properties for wastewater Treatment: A review. Bioresource technology, 126501.

Krishnan, K. A., & Keerthi, T. R. (2016). Comparative Evaluation of Bioactivities of two Marine Sponges, Zygomycale parishii and Callyspongia diffusa from Southwest Coast of India. British Journal of Pharmaceutical Research, 11(1).

Kwan, J. C., Meickle, T., Ladwa, D., Teplitski, M., Paul, V., & Luesch, H. (2011). Lyngbyoic acid, a “tagged” fatty acid from a marine cyanobacterium, disrupts quorum sensing in Pseudomonas aeruginosa. Molecular BioSystems, 7(4), 1205–1216.

Larimer, C., Winder, E., Jeters, R., Prowant, M., Nettleship, I., Addleman, R. S., & Bonheyo, G. T. (2015). A method for rapid quantitative assessment of biofilms with biomolecular staining and image analysis. Analytical and Bioanalytical Chemistry, 408(3), 999–1008. https://doi.org/10.1007/s00216-015-9195-z

Li, K. S., & Wah, C. S. (2017). Antioxidant and antibacterial activity of Acorus calamus. L leaf and rhizome extracts. Jurnal Gizi Klinik Indonesia, 13(4), 144–158.

Mannapperuma, W. M. G. C. K., Abaysekara, C. L., Herath, G. B. B., & Werellagama, D. R. I. B. (2013). Potentially pathogenic bacteria isolated from different tropical waters in Sri Lanka. Water Science and Technology: Water Supply, 13(6), 1463–1469.

Marasinghe, M. M. K. I. (2019). PhD dissertation “The Community Structure Dynamics and Invasive Potential of Biofouling Assemblage in Colombo Port”

Marasinghe, M. M. K. I., Ranatunga, R. R. M. K. P., & Anil, A. C. (2016). Bryozoans species composition in Colombo Port with a description of two new species. In Proceedings of International Forestry and Environment Symposium (Vol. 21).

Marasinghe, M. M. K. I., Ranatunga, R. R. M. K. P., & Anil, A. C. (2018). Settlement of non-native Watersipora subtorquata (d’Orbigny, 1852) in artificial collectors deployed in Colombo Port, Sri Lanka. Dr.lib.sjp.ac.lk. http://dr.lib.sjp.ac.lk/handle/123456789/6937

Marasinghe, M. M. K. I., Ranatunga, R. R. M. K. P., Anil, A. C., & Weerasinghe, R. L. (2015). Fitrst record of an invasive cheilostome bryozoan: Schizoporella errata in Sri Lankan waters. Proceedings of the 2^nd^ International Marine Symposium (pp. 23). Sri Lanka Foundation, Colombo, Sri Lanka.

Marine Environment Protection Authority-Sri Lanka. https://mepa.gov.lk/

Meenakshi, V. K., Veerabahu, C., & Roselin, K. F. (2013). GC-MS and IR Studies Of Ethanolic Extract of Colonial Ascidian-Polyclinum madrasensis Sebastian. 1952. International Journal of Pharma and Bio Sciences, 4(4), 1187–1198.

Menchaca, I., Zorita, I., Rodríguez-Ezpeleta, N., Erauskin, C., Erauskin, E., Liria, P., Mendibil, I., Santesteban, M., Urtizberea, I., (2014). Guide for the evaluation of biofouling formation in the marine environment. Revista de Investigación Marina, AZTI-Tecnalia, 21(4): 89–99

Mukherjee, P. K., Kumar, V., Mal, M., & Houghton, P. J. (2007). In vitro acetylcholinesterase inhibitory activity of the essential oil from Acorus calamus and its main constituents. Planta Medica, 73(3), 283–285.

Muzquiz, M. (2000). ALKALOIDS| Gas Chromatography.

National Center for Biotechnology Information. (2019, June 12). PubChem Compound Summary for CID 5281515, beta-Caryophyllene. PubChem. https://pubchem.ncbi.nlm.nih.gov/compound/beta-Caryophyllene

Ojala, J., & Tenold, S. (2017). Maritime trade and merchant shipping: The shipping/trade ratio since the 1870s. International Journal of Maritime History, 29(4), 838–854.

Peters, L., Wright, A. D., Krick, A., & König, G. M. (2004). Variation of Brominated Indoles and Terpenoids Within Single and Different Colonies of the Marine Bryozoan Flustra foliacea. Journal of Chemical Ecology, 30(6), 1165–1181. https://doi.org/10.1023/b:joec.0000030270.65594.f4

Ports & Terminals Insight. (2017). Drewry Maritime Research and Consulting Services. https://www.drewry.co.uk

Ranatunga, R. R. M. K. P., Asanthi, H. B., De Croos, M. D. S. T., Jayasiri, H. B., Kumara, P. B. T. P., Maithreepala, R. A., & Pathmalal, M. M. (2015). Port Biological Baseline Survey Report Volume I–Port of Colombo. Marine Environmental Protection Authority, Ministry of Environment and Mahaweli Development.

Richmond, M. D., & Seed, R. (1991). A review of marine macrofouling communities with special reference to animal fouling, Biofouling, 3:2, 151-168, DOI:10.1080/08927019109378169

Saeed, N. M., El-Demerdash, E., Abdel-Rahman, H. M., Algandaby, M. M., Al-Abbasi, F. A., & Abdel-Naim, A. B. (2012). Anti-inflammatory activity of methyl palmitate and ethyl palmitate in different experimental rat models. Toxicology and Applied Pharmacology, 264(1), 84–93.

Santalova, E. A., Makarieva, T. N., Gorshkova, I. A., Dmitrenok, A. S., Krasokhin, V. B., & Stonik, V. A. (2004). Sterols from six marine sponges. Biochemical Systematics and Ecology, 32(2), 153–167.

Schultz, M. P. (2007). Effects of coating roughness and biofouling on ship resistance and powering. Biofouling, 23(5), 331–341. https://doi.org/10.1080/08927010701461974

Schultz, M. P., Bendick, J. A., Holm, E. R., & Hertel, W. M. (2010). Economic impact of biofouling on a naval surface ship. Biofouling, 27(1), 87–98. https://doi.org/10.1080/08927014.2010.542809

Shaaban, M. T., Ghaly, M. F., & Fahmi, S. M. (2021). Antibacterial activities of hexadecanoic acid methyl ester and green-synthesized silver nanoparticles against multidrug-resistant bacteria. Journal of Basic Microbiology, 61(6), 557–568.

Si, S. C., Sree, A., Bapuji, M., Gupta, J. K., & Siddqui, K. I. (2001). Fatty acid composition of lipids of some of marine sponges from Orissa coast. Indian Journal of Pharmaceutical Sciences, 63(2), 158.

Sieniawska, E., Sawicki, R., Swatko-Ossor, M., Napiorkowska, A., Przekora, A., Ginalska, G., & Augustynowicz-Kopec, E. (2018). The effect of combining natural terpenes and antituberculous agents against reference and clinical Mycobacterium tuberculosis strains. Molecules (Basel, Switzerland), 23(1), 176.

Son, B. W., Kim, J. C., Lee, S. M., Cho, Y. J., Choi, J. S., Choi, H. D., & Song, J. C. (2000). New diacylgalactolipids from the marine cyanophycean microalga Oscillatoria sp. Bulletin of the Korean Chemical Society, 21(11), 1138–1140.

Sylvester, F., Kalaci, O., Leung, B., Lacoursière-Roussel, A., Murray, C. C., Choi, F. M., Bravo, M. A., Therriault, T. W., & MacIsaac, H. J. (2011). Hull fouling as an invasion vector: Can simple models explain a complex problem? Journal of Applied Ecology, 48(2), 415–423.

Takahashi, C., Kikuchi, N., Katou, N., Miki, T., Yanagida, F., & Umeda, M. (1995). Possible anti-tumour-promoting activity of components in Japanese soybean fermented food, Natto: Effect on gap junctional intercellular communication. Carcinogenesis, 16(3), 471–476.

Takamura, H., Iwamoto, K., Nakao, E., & Kadota, I. (2013). Total synthesis of two possible diastereomers of (+)-sarcophytonolide C and its structural elucidation. Organic Letters, 15(5), 1108–1111.

Todd, P. A., Heery, E. C., Loke, L. H., Thurstan, R. H., Kotze, D. J., & Swan, C. (2019). Towards an urban marine ecology: Characterizing the drivers, patterns and processes of marine ecosystems in coastal cities. Oikos, 128(9), 1215–1242.

Uebel, T., Hermes, L., Haupenthal, S., Müller, L., & Esselen, M. (2020). α-Asarone, β-asarone, and γ-asarone: Current status of toxicological evaluation. Journal of Applied Toxicology. https://doi.org/10.1002/jat.4112

Wahl, M. (1989). Marine epibiosis. I. Fouling and antifouling: some basic aspects. Marine ecology progress series, 58, 175–189.

Wesołowska, A., Grzeszczuk, M., Wilas, J., & Kulpa, D. (2016). Gas Chromatography-Mass Spectrometry (GC-MS) analysis of indole alkaloids isolated from Catharanthus roseus (L.) G. don cultivated conventionally and derived from in vitro cultures. Notulae Botanicae Horti Agrobotanici Cluj-Napoca, 44(1), 100–106.

WoRMS Editorial Board (2018). World Register of Marine Species. Available from https://www.marinespecies.orgat VLIZ. Accessed 2018-05-31. doi:10.14284/170

Wu, Z., Liu, D., Proksch, P., Guo, P., & Lin, W. (2014). Punctaporonins H–M: Caryophyllene-Type Sesquiterpenoids from the Sponge-Associated Fungus Hansfordia sinuosae. Marine Drugs, 12(7), 3904–3916. https://doi.org/10.3390/md12073904

Yusuf, A. Z., Zakir, A., Shemau, Z., Abdullahi, M., & Halima, S. A. (2014). Phytochemical analysis of the methanol leaves extract of Paullinia pinnata linn. Journal of Pharmacognosy and Phytotherapy, 6(2), 10–16. https://doi.org/10.5897/jpp2013.0299

