## Supplemental Table 1 for "Chemical diversity in some biofouling organisms from the western coastal waters of Sri Lanka"

Table S1 GC-MS analysis for chemical composition on the extracts of target organisms

| Chemical group | Chemical compound | Common name | Molecular formula | *Schizoporella errata* (a) | *Botrylloides violaceus* (b) | *Carijoa risei* (c) | *Acanthella cavernosa* (d) | *Clathria* sp. (e) | *Callyspongia diffusa* (f) | *Halichondria panicea* (g) | *Mycale parishii* (f) |
|---|---|---|---|---|---|---|---|---|---|---|---|
| Ester | Octadecanoic acid, ethyl ester | Ethyl stearate | $C_{20}H_{40}O_2$ | | | | | | + | | + |
| Fatty acid | Octadecanoic acid | Stearic acid | $C_{18}H_{36}O_2$ | + | | | | | | | |
| Ester | Tetradecanoic acid, ethyl ester | Ethyl tetradecanoate | $C_{26}H_{32}O_2$ | + | | | | | + | | |
| Fatty acid | Tetradecanoic acid | Myristic acid | $C_{14}H_{28}O_2$ | + | | | | | | | |
| Ester | Dodecanoic acid, methyl ester | Methyl laurate | $C_{13}H_{26}O_2$ | | + | | | | + | | + |
| Fatty acid | Dodecanoic acid | Lauric acid | $C_{12}H_{24}O_2$ | + | | | | | | | |
| Ester | Hexadecanoic acid, methyl ester | Methyl palmitate | $C_{17}H_{34}O_2$ | + | | + | + | | + | | + |
| Alkane | Eicosane | Icosane | $C_{20}H_{42}$ | | | | | + | | | + |
| Sterol | .gamma.-Sitosterol | Clionasterol | $C_{29}H_{50}O$ | | | | | + | | | |

| | | | | | | | |
|---|---|---|---|---|---|---|---|
| Sterol | Ergosta-5,22-dien-3-ol, (3β,22E) | Brassicasterol | $C_{28}H_{46}O$ | + | | | |
| Sesquiterpene | Caryophyllene | | $C_{15}H_{24}$ | | + | | + |
| Monoterpenoid | 1,6-Octadien-3-ol, 3,7-dimethyl | Linalool | $C_{10}H_{18}O$ | | + | | |
| Monoterpenoid | 2,6-Octadien-1-ol, 3,7-dimethyl-, (E) | Geraniol/Nerol | $C_{10}H_{18}O$ | | + | | |
| Sesquiterpene | Cyclohexane, 1-ethenyl-1-methyl-2,4-bis(1-methylethenyl)-, [1S(1.alpha.,2.beta.,4.beta.)] | .beta.-Elemene | $C_{15}H_{24}$ | | + | | |
| phenylpropanoid | Benzene, 1,2,4-trimethoxy-5-(1-propenyl)-, (Z) | Alpha-Asarone | $C_{12}H_{16}O_3$ | | + | | |
| Alkane | Hentriacontane | | $C_{31}H_{64}$ | | | + | |
